# Coordination games in cancer

**DOI:** 10.1101/2021.06.22.449436

**Authors:** Péter Bayer, Robert A. Gatenby, Patricia H. McDonald, Derek R. Duckett, Kateřina Staňková, Joel S. Brown

**Author notes:** We thank Ingela Alger, Dave Basanta, Monica Salvioli, and Jeffrey West, as well as seminar participants of Moffitt’s Integrated Mathematical Oncology Department and the Institute of Advance Studies in Toulouse for useful discussions and comments.

## Abstract

We propose a model of cancer initiation and progression where tumor growth is modulated by an evolutionary coordination game. Evolutionary games of cancer are widely used to model frequency-dependent cell interactions with the most studied games being the Prisoner’s Dilemma and public goods games. Coordination games, by their more obscure and less evocative nature, are left understudied, despite the fact that, as we argue, they offer great potential in understanding and treating cancer. In this paper we present the conditions under which coordination games between cancer cells evolve, we propose aspects of cancer that can be modeled as results of coordination games, and explore the ways through which coordination games of cancer can be exploited for therapy.

## 1 Introduction

Cancer cells engage in evolutionary games (Tomlinson, 1997; Gatenby and Vincent, 2003). For them to do so, they must exhibit two dynamics and certain types of interactions. As an ecological dynamic, cancer cells exhibit survival and proliferation giving rise to changes in their population sizes. As an evolutionary dynamic, the heritable traits of the population change as cancer clades with more successful phenotypes outcompete and replace those with less successful ones (Cunningham et al., 2011). To be an evolutionary game the success of a phenotype must be context dependent. Its success depends upon not just the number but the phenotypes of those cells with which it interacts. Thus, the “best” strategy for a cancer cell will depend upon the strategies of other cancer cells (Brown, 2016). Researchers have proposed and identified a number of games that may typically occur within a patient’s tumor. Cancer cells may engage in the Prisoner’s Dilemma (Kareva, 2011; West et al., 2016) and exhibit cooperation (Axelrod et al., 2006). In this case, a cancer cell by co-feeding neighboring cells or by secreting factors that improve the mirco-environment may incur a cost to itself while providing benefits to neighboring cancer cells. If the cooperative cancer cell also derives some benefit from its action then the game shifts from being a Prisoner’s Dilemma to a public goods game (Archetti and Pienta, 2019). By producing VEGF for recruiting vasculature or creating acidic conditions as immune-suppression, the focal cancer cell benefits itself while also benefiting neighbors (Kimmel et al., 2019; Nogales and Zazo, 2021). Such games can promote a diversity of cancer cell types, where some become producers while others act as free-loaders, contributing nothing to the public good (Bayer et al., 2018). Cancer cells may also engage in the Tragedy of the Commons. This happens when the cancer cells over-invest in metabolic pathways, transporters, and capacity so as to pre-empt other cancer cells from acquiring resources from a shared common pool. The interstitial fluids can be the commons, and it may be that cancer cells, for instance, over-express GLUT-1. The level of expression may be higher than would be optimal for the group, but it is advantageous for the individual cancer cell if it gains nutrients that otherwise would have been harvested by its neighbors (Bukkuri et al., 2021).

Game theory has been suggested as the framework for evolutionarily informed therapies where the physician aims to anticipate and steer the eco-evolutionary dynamics of the cancer towards better outcomes or outright cure (Gatenby et al., 2009). A clinical trial of castrate-resistant metastatic prostate modeled the cancer cells as having three possible strategies in a game that has similarities to a rock-scissors-paper game (Zhang et al., 2017). Androgen deprivation therapy (Lupron) eliminates cancer cells with one of the strategies (*T* + cells requiring endogenous testosterone) allowing a second strategy (*TP*, testosterone producing, cells) to over-proliferate. A second therapy (Abiraterone) eliminates both of these strategies allowing for a third strategy (*T* −, cells independent of testosterone) to proliferate and threaten the patient. By cycling abiraterone on and off in an adaptive fashion, the trial aimed to suppress *T* + and *TP* cells once tumors grew too large (therapy on) while shrinking tumors to a threshold (therapy off at this point) as a means of retaining *T* + and *TP* cells to competitively suppress *T* − cells Zhang et al. (2017); You et al. (2017); Cunningham et al. (2018).

One class of games that has not been explored in cancer are coordination games. Coordination games are characterized by incentive structures that reward conformity. Deviation by any player results in lower payoffs for the whole population. The most stringent subclass of coordination games is called *pure coordination* games. In these games, positive payoffs are only attainable if all individuals choose the same strategy, for any other strategy combination all individuals receive zero payoffs. A typical application of this game is the adoption a new technology standard. If all participants adopt the new technology, payoffs are maximized, if no one adopts the new technology payoffs are lower but positive, if some participants adopt while others do not, confusion follows and payoffs are zero (top left panel of Table 1). If the rewards of coordination are identical, the game is called *choosing sides*. A classic example of this game is driving on either side of the road. If participants all adhere to driving on the same side, the traffic flows and payoffs are positive, otherwise chaos ensues and payoffs are zero (top right panel of Table 1). A less stringent class in terms of punishing discoordination is called the *stag hunt* game. One strategy offers low rewards but does not require coordination, the other offers high rewards that can only be attained by coordination. The original version of the game features two hunters who decide on which animal to hunt, stag or hare: if both choose stag, they are able to acquire it through a joint effort for a large reward, if any hunter chooses hare, they are able to acquire it individually for a lower reward, but if one chooses stag while the other chooses hare, the stag hunter fails and receives a payoff of zero (bottom panel of Table 1).

**Table 1:**
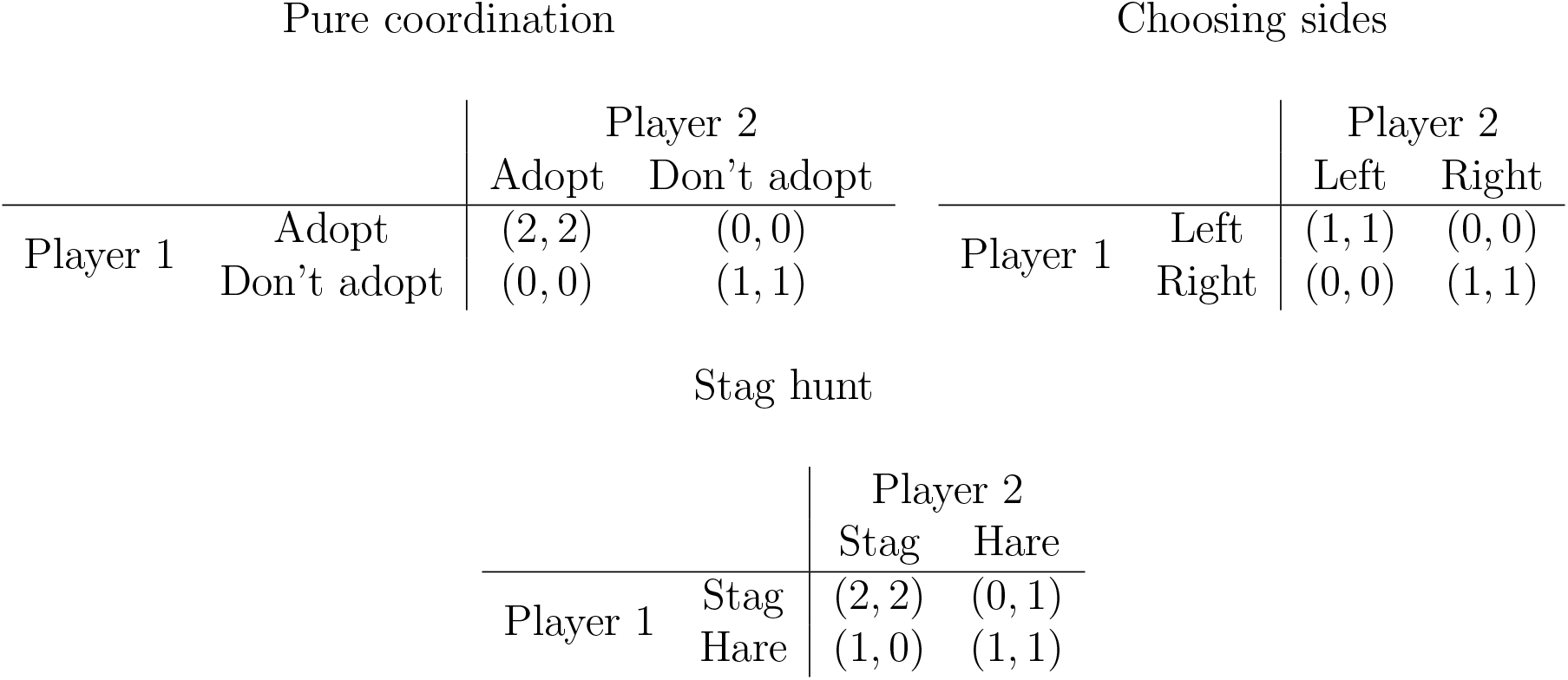
Typical payoff structures of classic coordination games.

Evolutionary coordination games replace the element of individual choice with heritable traits and payoffs with reproductive fitness. Populations playing evolutionary coordination games invariably develop similar traits as discoordinating individuals suffer fitness penalties. Furthermore, the populations that achieve coordination more quickly will thrive as a whole, while those that fail to coordinate may fail. Table 2 showcases a coordination game with generic payoff parameters. For a cancer cell, neither strategy is a good or bad one per se; rather its success depends on the predominant strategy in the population of cancer cells. If type 1 predominates, then a focal cancer cell does best being type 1 and vice versa when type 2 is predominant.

**Table 2:** An evolutionary coordination game with two strategies. Without loss of generality we assume *r*_1_ ≥ *r*_2_. To get a coordination game, we further assume *r*_1_ ≥ *d*_1_ and *r*_2_ ≥ *d*_2_, ensuring that each type proliferates in its own environment at a higher rate than in the other’s. We further assume *r*_2_ ≥ *d*_1_, ensuring that 2 remains a viable strategy in its own environment and cannot be invaded by 1 (*r*_1_ ≥ *d*_2_ follows from the previous assumptions, ensuring that 1 cannot be invaded by 2).

|  |  | Predominant strategy |  |
| --- | --- | --- | --- |
|  |  | Type 1 | Type 2 |
| Focal cell | Type 1 | $r_1$ | $d_1$ |
| | Type 2 | $d_2$ | $r_2$ |

This makes identifying coordination games in cancer challenging as the two competing types should not be found together. Their co-occurrence within the same tumor microenvironment or even within the same tumor or patient seems unlikely and would manifest only as a transient dynamics as either type 1 or 2 come to predominate. Furthermore, the predominance of type 1 in one part of a tumor and that of type 2 in another is not strong evidence for a coordination game. For instance, the edge of a tumor may favor 1 and the interior 2 independent of which cell type is initially predominant in that region (Lloyd et al., 2016).

In a coordination game it is the predominance of type 1 at the site that makes type 2 unsuitable. Had 2 become predominate first then 1 would be absent. It is this priority effect that characterizes the coordination game in cancer. Whichever strategy gets established first among the cancer cells excludes the other. This makes it difficult to spot and identify coordination games by observing the outcome within an individual patient. The alternative strategies that the population *could have* coordinated upon are no longer visible. If coordination games exist in cancer, by the time the disease is detected, most cancer patients would likely present with a cancer that has already evolved to a common phenotype. As a result, the various strategies of these games are more likely to manifest *between* rather than *within* patients.

The structure of the paper is as follows. In Section 2 we raise examples of possible coordination games in cancer. In Section 3 we build a Lotka-Volterra competition model of cancer where growth is regulated by an underlying coordination game. In Section 4 we investigate the interaction between coordination games of cancer growth and resistance to cytotoxic therapy in a sensitive-resistant model and showcase how therapies that take into account the coordination game can lead to better outcomes. In Section 5 we showcase how therapy outcomes can be improved further by therapy that increases the transmutation rates between the two competing phenotypes. Section 6 contains the concluding discussion. An brief appendix shows the mathematical conditions and basic properties of coordination games.

## 2 Possible coordination games in cancer

The concept of “driver genes” may help identify a coordination game. In many cancers, a specific gene mutation is observed in virtually all cells and is thought to be a critical event that provides a steady oncogenic signal to maintain survival and proliferation of the cancer cells. Perhaps the best example of this is the oncogenic mutation of Epidermal Growth Factor in lung cancers. This phenotype accounts for 15-20% of clinical lung cancers. Unlike most lung cancer cohorts in which prolonged smoking and exposure to air pollutants are clear risk factors, EGFR-mut cancers tend to occur in younger patients who are non-smokers or have had limited exposure. Typically, EGFR-mut lung cancer cohorts tends to be young, Asian, and female. For these patients, the EGFR-mutation occurs in essentially all of the cancer cells of the primary and metastatic tumors (Yatabe et al., 2011). Furthermore, the overall mutational burden in EGFR-mut lung cancer is significantly smaller than EGFR WT lung cancers, indicating a molecularly more homogeneous intra-tumoral population (Jiao et al., 2019). We propose that EGFR-mut versus EGFR WT lung cancers represent a coordination game.

Treatment of EGFR-mut lung cancers with tyrosine kinase inhibitors (TKI) that specifically target the EGFR-mut function results in a complete or partial response in about 75% of patients (Shim et al., 2013). However, treatment response is transient and tumor progression occurs within 12 to 14 months. By molecular analysis, at least 7 different strategies permit the lung cancer cells to overcome the TKIs (mechanisms in 15% of cases remain unknown). Furthermore, second line treatments with chemotherapy or immunotherapy show minimal efficacy (Lee et al., 2017). It has been noted that in many cases, resistance takes the form of the cancer cells losing the EGFR-mut and becoming like EGFR-WT lung cancers. As a coordination game, treating the EGFR-mut may simply shift the cancer to an alternate stable state.

Another subset of lung cancers (about 40%) have an oncogenic KRAS mutations indicating that driver mutations are, to some extent substitutable. But cancer cells with different driver mutations may not be able to coexist within the same patient or tumor. In the case of non-small-cell lung cancer, for example, the driver mutations in KRAS and EGFR seem to be mutually exclusive suggesting that, once a cancer population has one, the other is actively selected against. Thus, one or the other is beneficial, but both are deleterious to the cancer cell. This is necessary for a coordination game but not sufficient. While exhibiting both mutations in a single cell may be selected against, it does not mean that an EGFR mutant cancer cell would be at a disadvantage in a community of KRAS-mutant cancer cells and vice-versa. For these two strategies to be a coordination game, a particular cancer cell with a particular driver mutation would have to be more successful if its neighbors harbored the same driver mutation.

In patients with breast cancer, the ubiquitously expressed beta-arrestin isoforms (*β*-arrestin 1; ARRB2, and *β*-arrestin 2; ARRB3) may form the biological basis for a coordination game. Beta-arrestins function as “terminators” of G protein-coupled receptor (GPCR) signaling. More recently, *β*-arrestins, by virtue of their scaffolding functionality, have also been shown to serve as signal “transducers”. As such, *β*-arrestins play roles in MAPK signaling (McDonald and Lefkowitz, 2001), and the regulation of several basic cellular functions including cell cycle regulation (Cao et al., 2017), proliferation, cell migration (Bostanabad et al., 2021), apoptosis (Kook et al., 2014), and DNA damage repair (Hara et al., 2011; Nieto et al., 2020). Data supporting a physiologically relevant role for these *β*-arrestin-mediated responses are nowhere more compelling than in cancer.

Multiple independent studies have reported a change in the expression of *β*-arrestins in breast cancer cells and patient tumors, wherein changes in the expression of *β*-arrestins correlate with poor patient survival (Michal et al., 2011), with *β*-arrestin 2 expression serving as a prognostic biomarker in the clinical course of breast cancer. Investigations of several human genomic datasets revealed that the expression of *β*-arrestin 1 is downregulated in triple negative breast cancer, the most aggressive breast cancer subtype in terms of poor outcomes and high rates of relapse (Son et al., 2019). This suggests that cancer cell fitness is maximized by changing the expression of one but not both *β*-arrestins. Changing one promotes self-sufficiency in proliferative signaling, while keeping the other *β*-arrestin unchanged maintains necessary basic cellular functions.

While these observations can explain how a co-adapted set of genes coordinates cellular processes within a cancer cell, it does not necessarily qualify as a coordination game for explaining why all cells of the patient’s cancer show one pattern of changes in *β*-arrestin and not the other. The coordination game may result from the way *β*-arrestins control aspects of cell-cell signaling through the regulation of GPCR activity, as well as other cell surface receptors. In this case, the value of a particular isoform of *β*-arrestin for successful cell-cell communication within the tumor may be determined by the unified expression of this isoform by surrounding cancer cells. That is, the dominant *β*-arrestin isoform may define rules of the road for all of the cancer cells.

## 3 Coordination game and cancer growth dynamics

We propose a two-phenotype model of cancer growth with cell types 1 and 2 with the payoffs *r*_1_*, r*_2_*, d*_1_*, d*_2_ satisfying the conditions of a coordination game (Table 2).

Our eco-evolutionary setting is described by a pair of ordinary differential equations (ODEs), modeling the population of the two types. Our model closely resembles Lotka-Volterra competition equations with the exception that tumor growth depends endogenously on its composition. The model features a joint carrying capacity (*K*) and joint strong Allee-effect (with Allee-threshold *T*) for the cell types. We add type-specific death rates (*c*_1_*, c*_2_) to capture the immune system’s function of destroying cancer cells and transmutation rates (*m*_1_*, m*_2_) capturing cancer’s mutation one type to the other. Finally, we account for possible non-linearities of fitness with respect to cell-type frequencies (with convexity parameter *α* ≥ 1) in order to capture a more general form of frequency-dependent growth. The model is summarized in Table 3.

**Table 3:** The ingredients and equations of describing a model of growth governed by a coordination game.

| Variables |  | Parameters |  |
| --- | --- | --- | --- |
| $x_1(t)$ | Type 1 population | $K$ | Carrying capacity |
| $x_2(t)$ | Type 2 population | $T$ | Allee-threshold |
| $x(t)$ | Tumor size ($x_1(t) + x_2(t)$) | $c_1, c_2$ | Cell death rates |
| $g(t)$ | Tumor composition ($x_1(t)/x(t)$) | $m_1, m_2$ | Mutation rates |
| | | $\alpha$ | Convexity parameter |

For tumors exhibiting coordination games, the cancer cells’ proliferation rates will be higher if the cellular composition is homogeneous. Furthermore, as the tumor grows its composition will converge on one or the other cell types as the predominant type. Both of these effects become stronger with a larger value for the convexity parameter. As the convexity parameter becomes larger so too does the punishment for discoordination. The mutation rate has the opposite effect. Increasing the mutation rate diversifies the composition of cancer cell types, reduces the fitness benefits achievable from coordination, and reduces the fitness of all of the cancer cells.

With respect to the composition of cancer cell, there exist two peaks of tumor fitness one at purely type 1 and the other at purely type 2. The extinction thresholds (because of the Allee effect) are lower and the maximum tumor sizes are higher when the tumor is homogeneous for one cell type or the other than when the tumor is heterogeneous (Figure 1).

**Figure 1:**
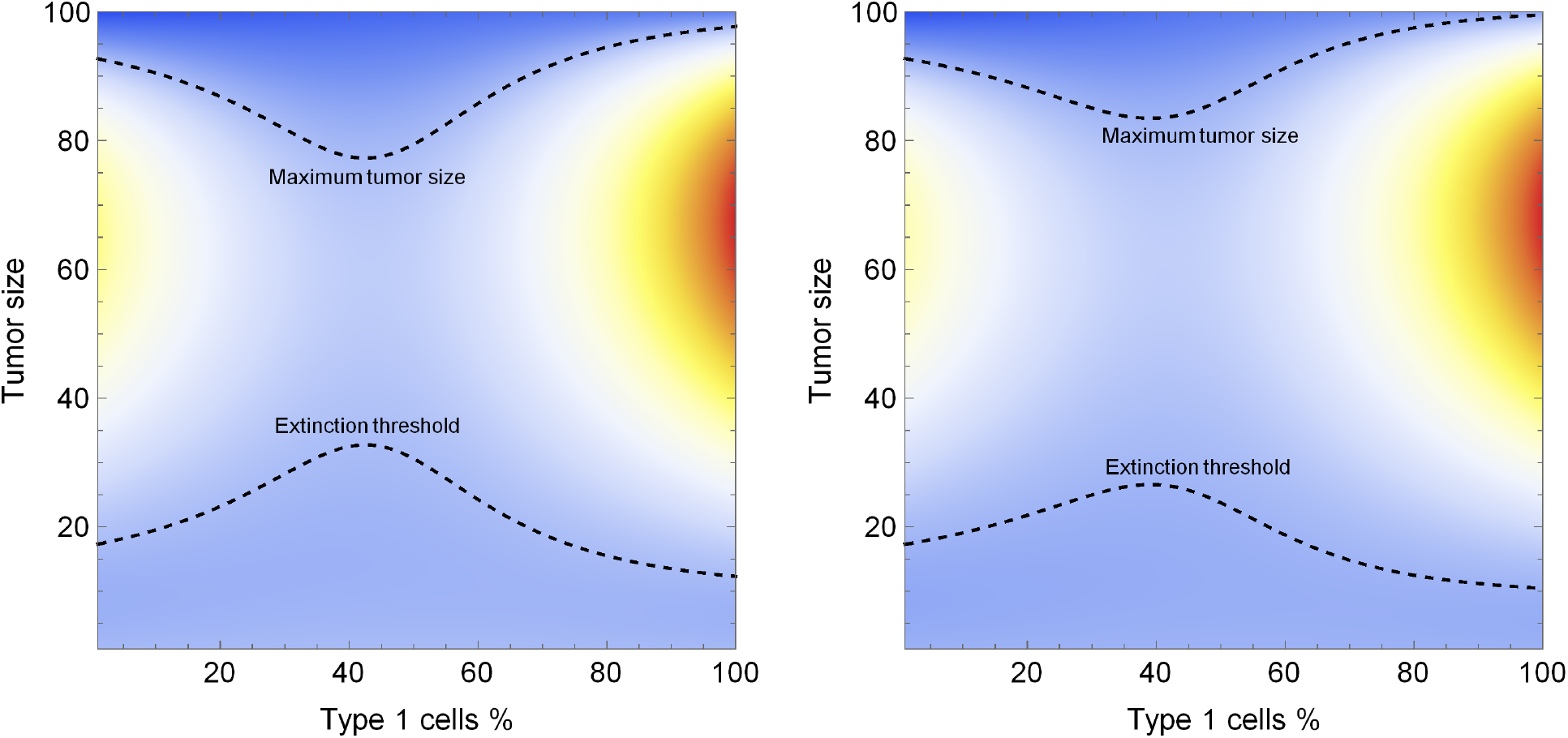
Heat map of tumor growth under a convex pure coordination game. Type 1’s growth rate is larger and its death rate is lower, therefore its fitness peak is higher, but a pure type 2 tumor also has a fitness peak. Decreasing type 1’s death rate lowers its extinction threshold and increases its maximum tumor size. Parameters: *r*_1_ = 0.25, *r*_2_ = 0.17, *d*_1_ = *d*_2_ = 0, *K* = 100, *T* = 10, *α* = 2, *c*_2_ = 0.1, left panel: *c*_1_ = 0.05, right panel: *c*_1_ = 0.01.

Provided that the mutation rates are sufficiently small, a tumor that does not go extinct will achieve coordination on one or the other cancer cell type. We highlight two typical calibrations of parameters. The two cancer cells types may be relatively symmetric in the sense that each can promotes similar tumor growth rates and sizes when growing in a monotypic tumor. If cancer cell types are close to symmetric in this sense, then the tumor outside the extinction zone may converge to one of two stable equilibria, with either type 1 or type 2 becoming the predominant phenotype. There also exist four unstable equilibria: one for each phenotype on the border of the extinction zone and two mixed equilibria, one on the border of the extinction zone, and one with a high tumor burden (Figure 2, left panel).

**Figure 2:**
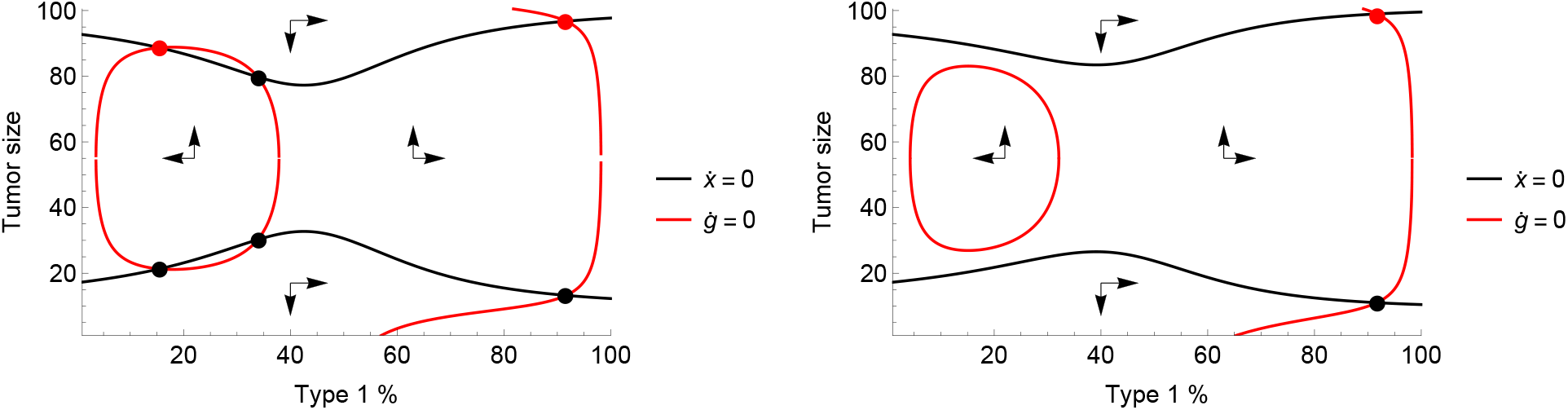
The phase diagram of tumor composition and size. On the left panel there are two stable equilibria (red dots) and two corressponding basins of attraction for tumors above the extinction zone: either coordination is achieved on type 1, or type 2. On the right panel there is a single stable equilibrium comprising mostly type 1 cells. Black dots denote unstable equilibria. Parameters: *r*_1_ = 0.25, *r*_2_ = 0.17, *d*_1_ = *d*_2_ = 0, *K* = 100, *T* = 10, *α* = 2, *m*_1_ = *m*_2_ = 0.01, *c*_2_ = 0.1, left panel: *c*_1_ = 0.05, right panel: *c*_1_ = 0.01.

If the cell types are sufficiently asymmetric in the sense that one of them provides a much higher tumor growth rate and/or lower death rate, then there only exists a single stable equilibrium. This equilibrium sees the cancer cell with the higher intrinsic growth rate or lower death rate becoming the predominant type with all growth trajectories pointing towards it (Figure 2 right panel). There is also one unstable equilibrium for this cancer cell type on the border of the extinction threshold.

## 4 Coordination games and cytotoxic therapies

Coordination games of cancer can be leveraged to improve the outcomes of existing cancer therapies. The challenge with many forms of cancer therapies such as chemotherapies is the evolution of resistance. Introducing a cytotoxic agent to attack the tumor yields a good initial response, with each subsequent use of the drug producing diminishing returns until the onset of resistance at which point the drug is ineffective.

In this section we model a general class of cytotoxic therapies and the onset of resistance as two strategies of a coordination game. Type 1 is *sensitive* to therapy, type 2 is *resistant*. We model the type’s response to therapy by functions (*γ*_1_(*t*), *γ*_2_(*t*)) which add to the cells’ death rates when the patient is receiving therapy. Compared to the base model of Section 1, we also omit the Allee-effect to better showcase the effect of leveraging the coordination game in eliminating the tumor. The model equations are then

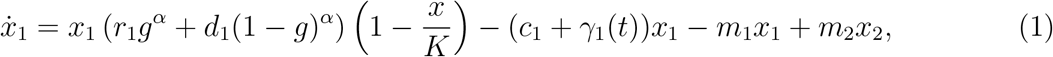

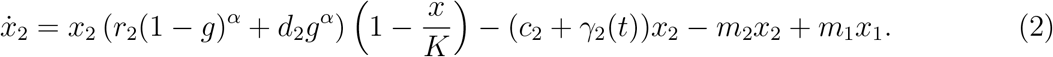

For *i* = 1, 2 and we have *γ_i_*(*t*) = *γ_i_* for time intervals when therapy is administered, indicating a constant dosage, with *γ*_1_ > *γ*_2_ ≥ 0 and *γ_i_*(*t*) = 0 when therapy is not administered. In the most extreme case, we have *γ*_2_ = 0 indicating that type 2 is completely unaffected by the cytotoxic therapy. We illustrate the system in Figure 3.

**Figure 3:**
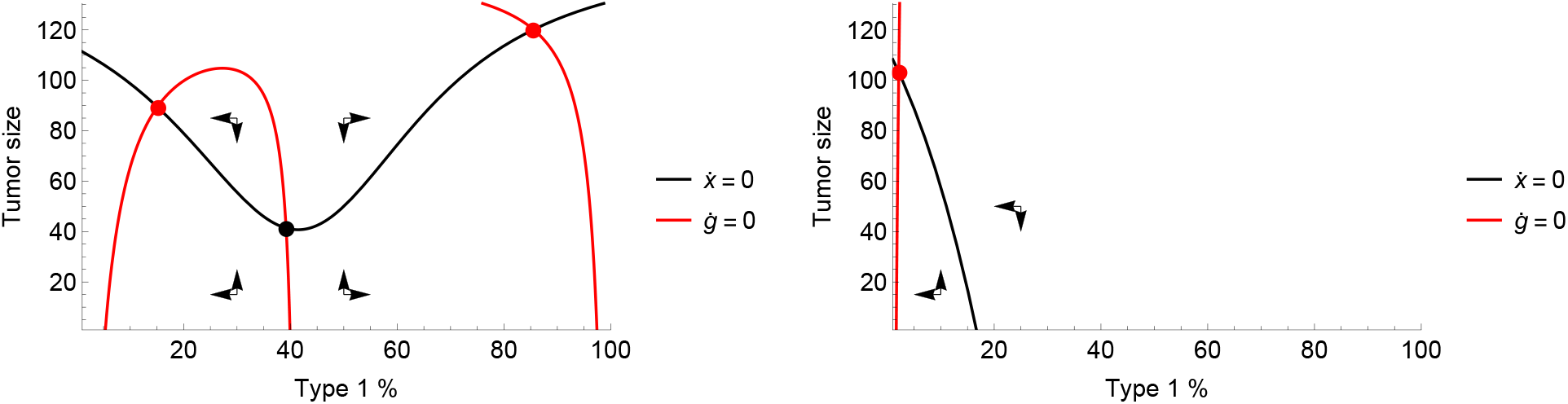
The zero-isoclines of the system defined by (1)-(2) with therapy off (left) and on (right). If therapy is off, there are two stable equilibria (red dots) comprising mostly type 1 and mostly type 2 cells, respectively, and a mixed unstable equilibrium (black dot). With therapy on, the only stable equilibrium consists of mostly type 2 cells. Parameters: *r*_1_ = 0.4, *r*_2_ = 0.2, *d*_1_ = *d*_2_ = 0, *K* = 150, *α* = 2, *c*_1_ = *c*_2_ = 0.05, *m*_1_ = *m*_2_ = 0.01, *γ*_1_ = 0.4, *γ*_2_ = 0.

Resistance occurs when an initially sensitive (type 1) tumor is disrupted by the therapy and achieves coordination on the resistant strategy (type 2). Upon initiation of therapy, the tumor size is reduced as type 1 cells are killed off. The tumor’s composition becomes more and more mixed, further lowering its fitness and reinforcing the therapy. If therapy continues uninterrupted, the resistant type 2 cells will eventually come to predominate in the tumor and, in a competitive release from type 1 cells, the tumor population returns to high levels (top left panel of Figure 4). Therapy may be turned off in hopes of avoiding the basin of attraction of the resistant type. For the parameters considered by Figure 3, this is reached when the frequency of type 1 cells drops below 40%. If therapy is turned off after this point, the tumor cannot be stopped from reaching an equilibrium of mostly type 2 cells (top right of Figure 4), if therapy is turned off any time before that, it will return to the stable equilibrium with mostly type 1 cells where therapy can once again be effective (bottom left of Figure 4).

**Figure 4:**
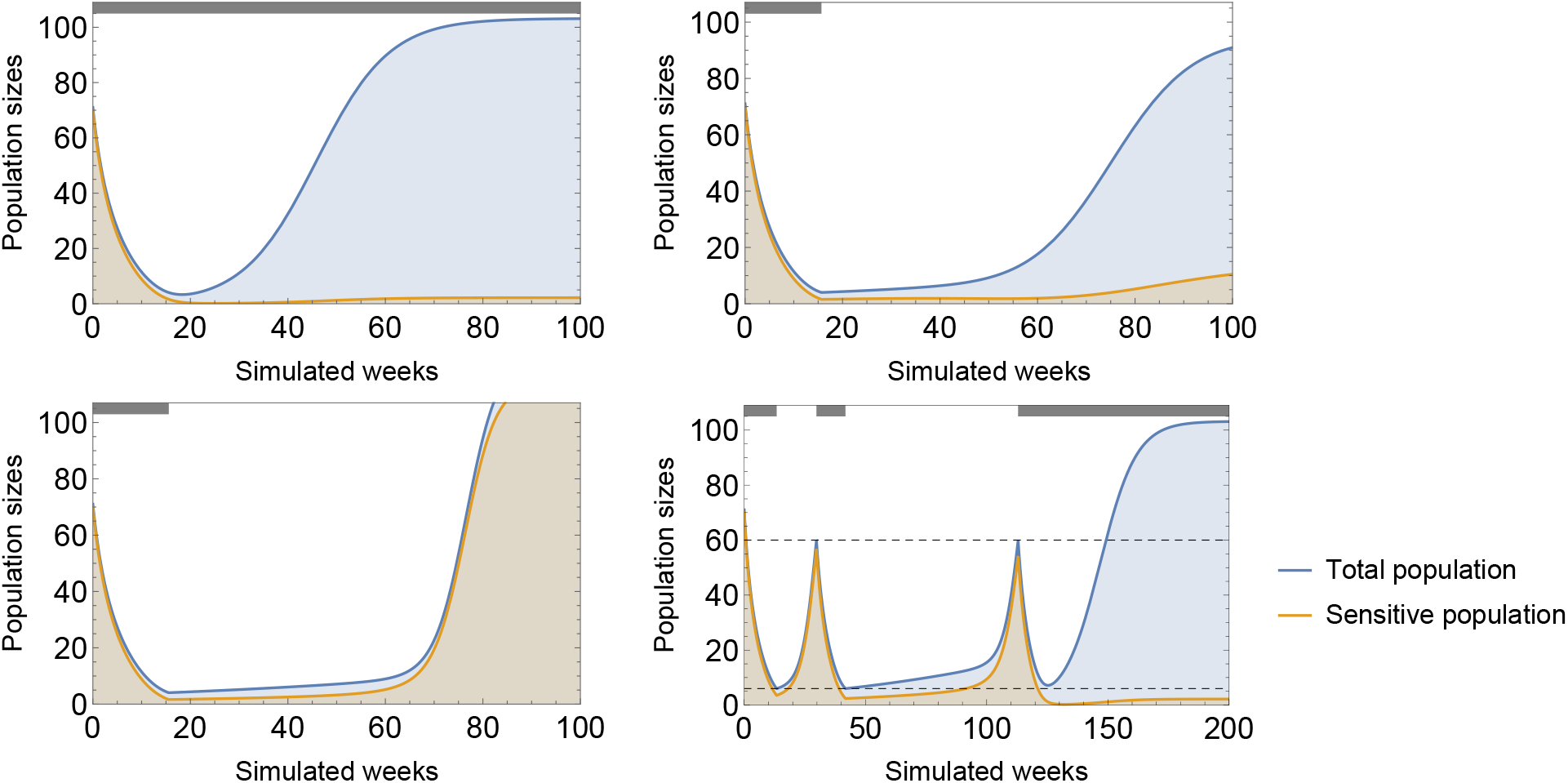
Continuous cytotoxic therapy yields a good initial response, drives down tumor burden, but the population rebounds once cancer coordinates on the resistant type (top left). If therapy is stopped at *g* = 39%, then by that time the tumor has entered the resistant type’s basin of attraction (top right). If therapy is stopped at *g* = 40%, the tumor remains in the sensitive type’s basin of attraction (bottom left). Under adaptive therapy control over the tumor is lost when it enters the resistant basin of attraction, which in the present case is after 3 treatment cycles (bottom right). Parameters: *r*_1_ = 0.4, *r*_2_ = 0.2, *d_a_* = *d_b_* = 0, *K* = 150, *c_a_* = *c_b_* = 0.05, *m_a_* = *m_b_* = 0.01, *α* = 2, *γ*_1_ = 0.4, *γ*_2_ = 0.

Adaptive therapies of cancer seek to retain control over the tumor by stopping therapy before resistance is complete, and restarting therapy only when the tumor burden becomes threatening. In a coordination game, adaptive therapy must strike a balancing between staying within the sensitive type’s basin of attraction to maintain control, and avoiding dangerous levels of the tumor burden. In the bottom right panel of Figure 4, the cytotoxic therapy is turned off when the tumor burden falls below 6 and turned on again when it exceeds 60. Control over the tumor is maintained for three on-off cycles.

Making use of the coordination game can improve therapy outcomes. As seen in Figure 4, mixed tumors have a slow rate of growth even when the tumor is small. Therapy regimens can be designed to keep the tumor composition mixed as follows: have cytotoxic therapy on until the tumor composition reaches a lower bound, 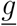, such that the tumor does not leave the sensitive type’s basin of attraction. Upon reaching the lower bound, therapy is turned off, until composition reaches an upper bound, 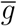. Figure 5 shows that such therapies, if well calibrated, can lead to the extinction of the tumor.

**Figure 5:**
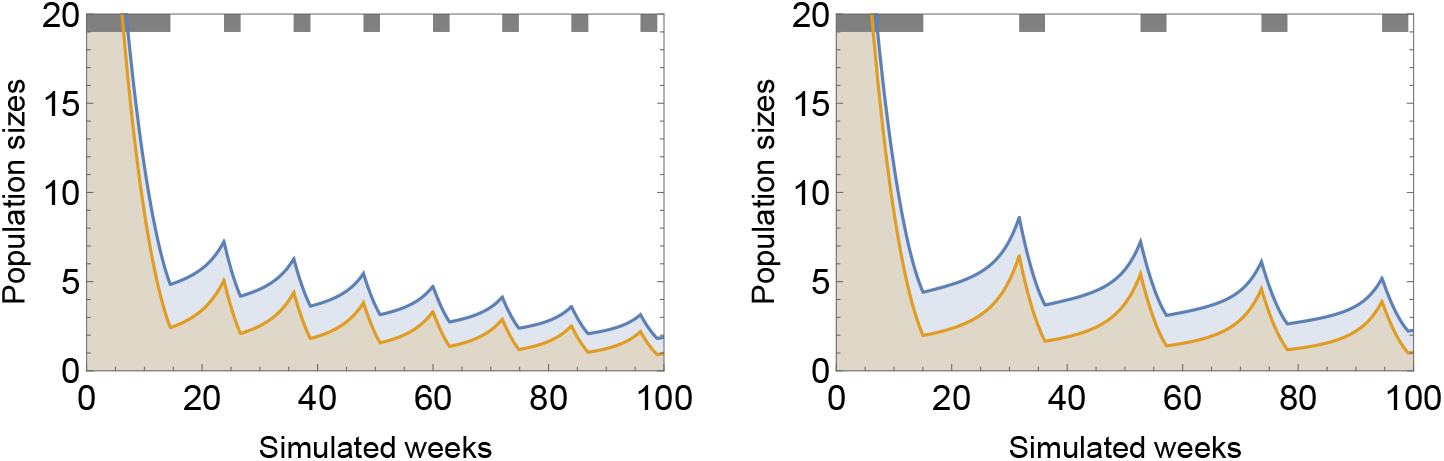
Cytotoxic therapy conditioned on the tumor’s composition. Therapy is turned off when *g*(*t*) falls below the lower bound 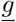 and it is turned on again when it rises above the upper bound 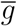. Both therapy regimens are able drive the tumor to extinction. A larger range between the bounds produces much a lower frequency of the treatment cycles. Parameters: *r*_1_ = 0.4, *r*_2_ = 0.2, *d_a_* = *d_b_* = 0, *K* = 150, *c_a_* = *c_b_* = 0.05, *m_a_* = *m_b_* = 0.01, *α* = 2, *γ*_1_ = 0.4, *γ*_2_ = 0, left panel: 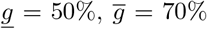, right panel: 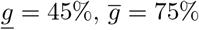.

Practically, conditioning treatment on the tumor’s composition is highly problematic as the prevalence of resistance can only be estimated from data on the tumor’s response to the cytotoxic therapy. In case of adaptive therapy in practice, decisions on stopping or restarting therapy are based on overall tumor size (using tumor biomarkers as proxies). Treatment regimens conditioned on tumor size are, predictably, much less reliable in controlling both the tumor size and its composition and therefore produce more modest results. The strategy is similar as before: apply therapy and until tumor size reaches a lower bound 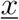, stop, then restart when tumor size exceeds an upper bound 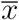.

The lower bound cannot be too low so as not to push the tumor into the resistant basin of attraction, while the upper bound needs to be high enough as to give the sensitive population enough time to replenish. The difference between the two bounds defines the *amplitude* of the treatment.

While such therapies are not precise enough to keep tumor at a mixed composition and thus achieve tumor extinction, a broad range of treatment strategies exist that put the tumor’s size and composition in a persistent and sustainable cycle, thus the tumor could be kept under control indefinitely. In Figure 6 we consider four adaptive treatment plans in convex coordination games, one high-amplitude, and three low-amplitude therapies at a small population, medium population, and large population, respectively. Each of these maintains control over the tumor indefinitely.

**Figure 6:**
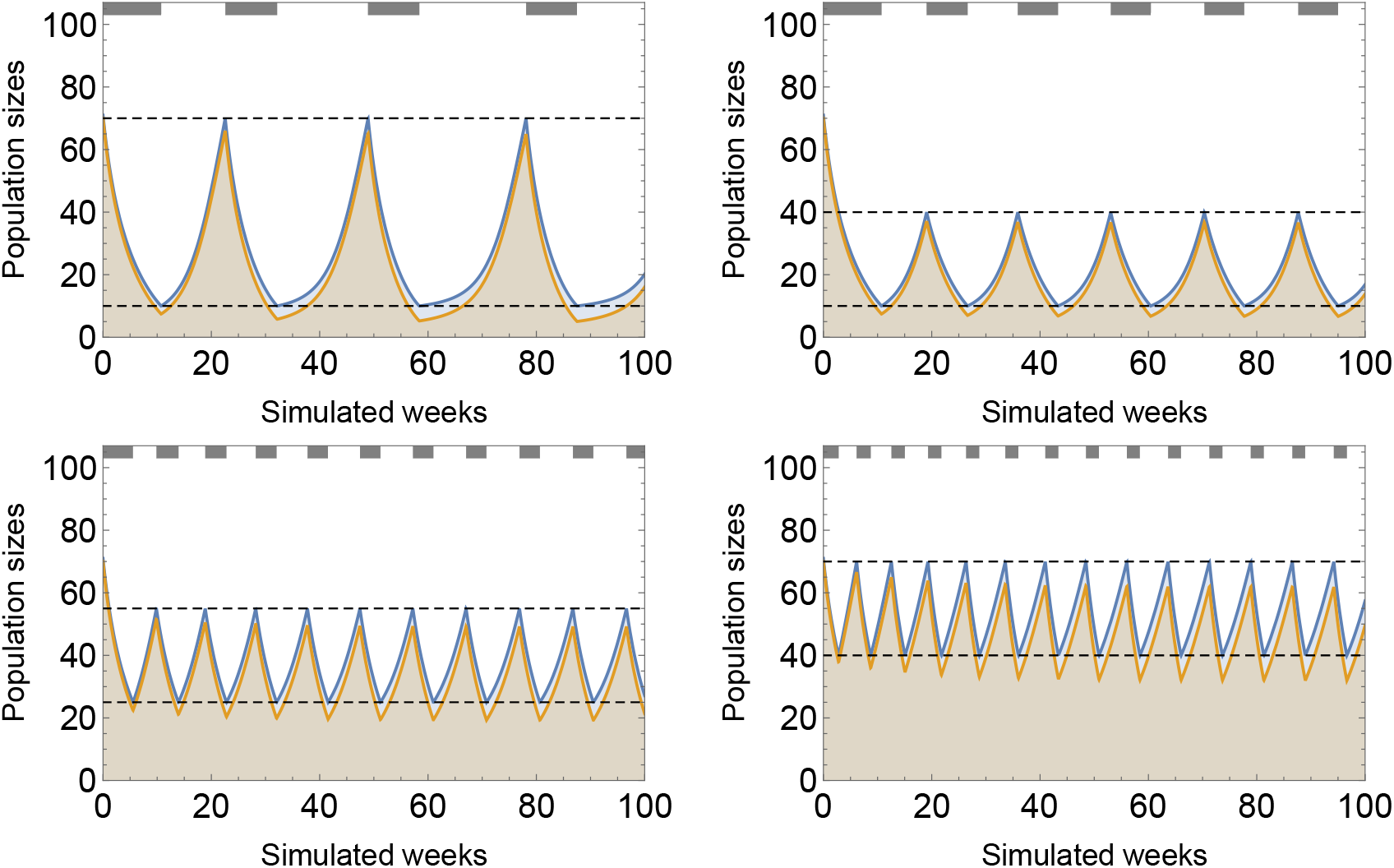
Four adaptive therapy strategies: *high amplitude* (top left), *low amplitude at small population* (top right left), *low amplitude at medium population* (bottom left), and *low amplitude at large population*. The tumor is kept under control indefinitely in each one. Parameters: *r*_1_ = 0.4, *r*_2_ = 0.2, *d_a_* = *d_b_* = 0, *K* = 150, *c_a_* = *c_b_* = 0.05, *m_a_* = *m_b_* = 0.01, *α* = 2, *γ*_1_ = 0.4, *γ*_2_ = 0.

In our example, each plan of adaptive therapy is able to keep control of the tumor and each offers a different set of advantages an disadvantages. Setting the lower bound at a small population produces a therapy with longer treatment cycles. Practically, this is beneficial but a low lower bound also means the tumor composition approaches the resistant type’s basin of attraction, risking losing control over the tumor. Setting a high upper bound gives the tumor more time to recover its composition which lowers the risk of losing control but exposes the patient to periods of high tumor burden. For a given amplitude of on-off cycling, deciding between a planned therapy aimed at maintaining small, medium, or a large population results in a tradeoff between the frequency of cycles, the risk of losing control, and periods of high tumor burden. In Table 4 we report these tradeoffs explicitly for the four adaptive treatment strategies considered in Figure 6.

**Table 4:** The characteristics of the four adaptive treatment strategies considered in Figure 6. High-amplitude adaptive therapy produces long cycles at the cost of approaching the resistant type’s basin of attraction and periods of high tumor burden. Low-amplitude therapies trade off the length of treatment cycles, which are the longest at small population levels, against the risk of losing control, which is lowest at large population level.

| Treatment strategy $(x, \bar{x})$ | Length of cycles<br>(Sim. weeks) | Maximum<br>tumor burden | Minimum<br>composition (%) |
| --- | --- | --- | --- |
| High-amplitude (10, 70) | 31 | 70 | 49 |
| Low-amplitude, small pop. (10, 40) | 17 | 40 | 66 |
| Low-amplitude, medium pop. (25, 55) | 10 | 55 | 77 |
| Low-amplitude, large pop. (40, 70) | 8 | 70 | 80 |

## 5 Mutation-promoting therapy

As shown by the previous section, coordination games may be leveraged for improving cytotoxic therapy outcomes. In the extreme, adaptive therapy in the context of a coordination game can engineer and extinction following a certain number of on-off cycles of the drug. More practically, on-off adaptive therapy strategies can indefinitely maintain a non-lethal tumor burden with a controllable composition. Up until this point, the only tool in the simulated physician’s arsenal was cytotoxic therapy, i.e. increasing the sensitive cell type’s death rate.

A more straightforward method for taking advantage of the coordination game involves lowering tumor fitness by maintaining a mixed composition of cancer cell types. One way to do this is by promoting mutation between the two types. We model this by adding type-specific response functions (*σ*_1_(*t*)*, σ*_2_(*t*)) which add to the cells’ mutation rates while the patient is under this type of therapy, called mutation therapy. Together with cytotoxic therapy, the model equations become

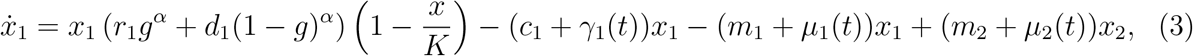

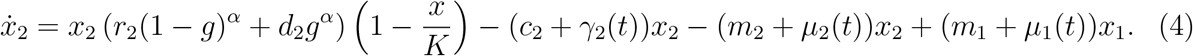

For *i* = 1, 2 and we have *μ_i_*(*t*) = *μ_i_* for time intervals under mutation therapy, with *μ*_1_*, μ*_2_ ≥ 0, and *μ_i_*(*t*) = 0 otherwise. Crucially, we allow both types’ mutation therapies, and cytotoxic therapy to be administered at different times, i.e. at any time the patient may receive any combination of the three, or may receive no therapy at all.

The role of mutation therapy is not to attack the tumor directly, but rather, to achieve or maintain a heterogeneous composition of cell types so that the tumor’s growth is inhibited by the mechanics of the coordination game, and direct attacks launched by other means are more effective. Mutation therapy pushes the stable composition isocline curve inwards, thus the stable equilibria will be more heterogeneous. If mutation therapy is strong enough, it may eliminate one of the stable equilibria, forcing the tumor to coordinate on a selected type. In Figure 7 we showcase the effects of mutation therapy in the system defined by (3)-(4).

**Figure 7:**
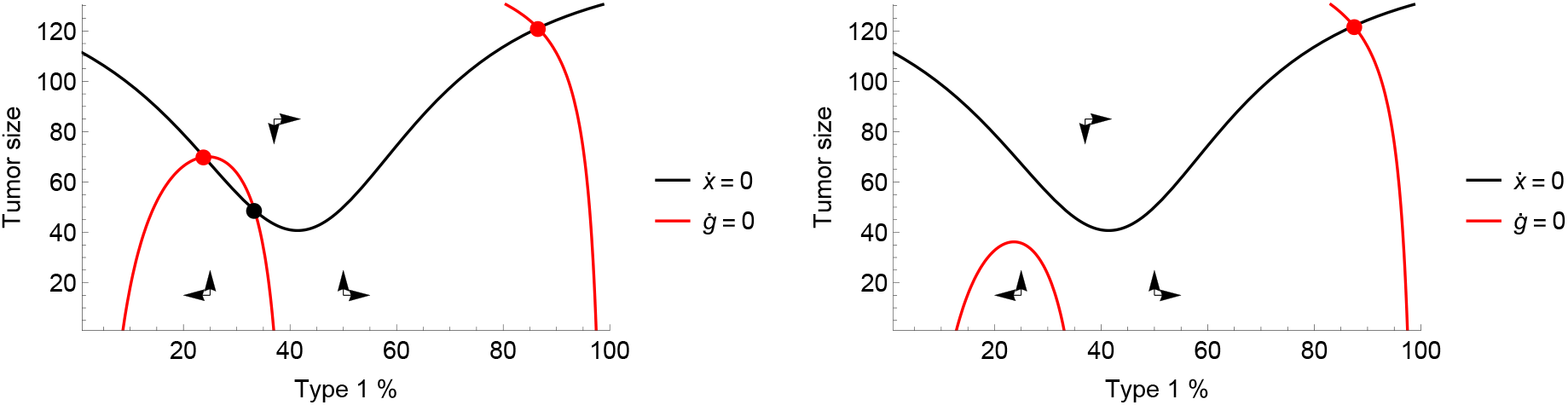
The effects of mutation therapy. Mutating type 2 cells to type 1 cells shrinks the former’s basin of attraction. A slight mutation therapy maintains two stable equilibria, one for each type (left), a stronger one eliminates the type 2-dominated stable equilibrium (right). Parameters: *r*_1_ = 0.4, *r*_2_ = 0.2, *d_a_* = *d_b_* = 0, *K* = 150, *c_a_* = *c_b_* = 0.05, *m_a_* = *m_b_* = 0.01, *α* = 2, *μ*_1_ = 0, left: *μ*_2_ = 0.005, right: *μ* = 0.01.

In the sensitive-resistant model, this effect can be exploited to force coordination on the sensitive type by mutating the resistant type to a sensitive one. We identify three ways by which doing so would improve therapy outcomes. First, applying continuous cytotoxic therapy together with mutation therapy delays coordination on the resistant type resulting in a longer progression time than without mutation therapy. Second, applying cytotoxic therapy adaptively is made safer with mutation therapy as the resistant type’s basin of attraction becomes smaller, thus the risk of losing control is greatly reduced. Third, new types of therapy aimed at the extinction of the tumor become possible. Figure 8 shows each of these in turn: (1) mutation therapy delays tumor progression under continuous cytotoxic therapy (top left panel), (2) mutation therapy makes it possible for a successful adaptive therapy to reach as low as *g* = 32% without losing control over the tumor (top right), and (3) mutation therapy makes it possible to engineer a successful extinction therapy (bottom right) that otherwise would be impossible in its absence (bottom left).

**Figure 8:**
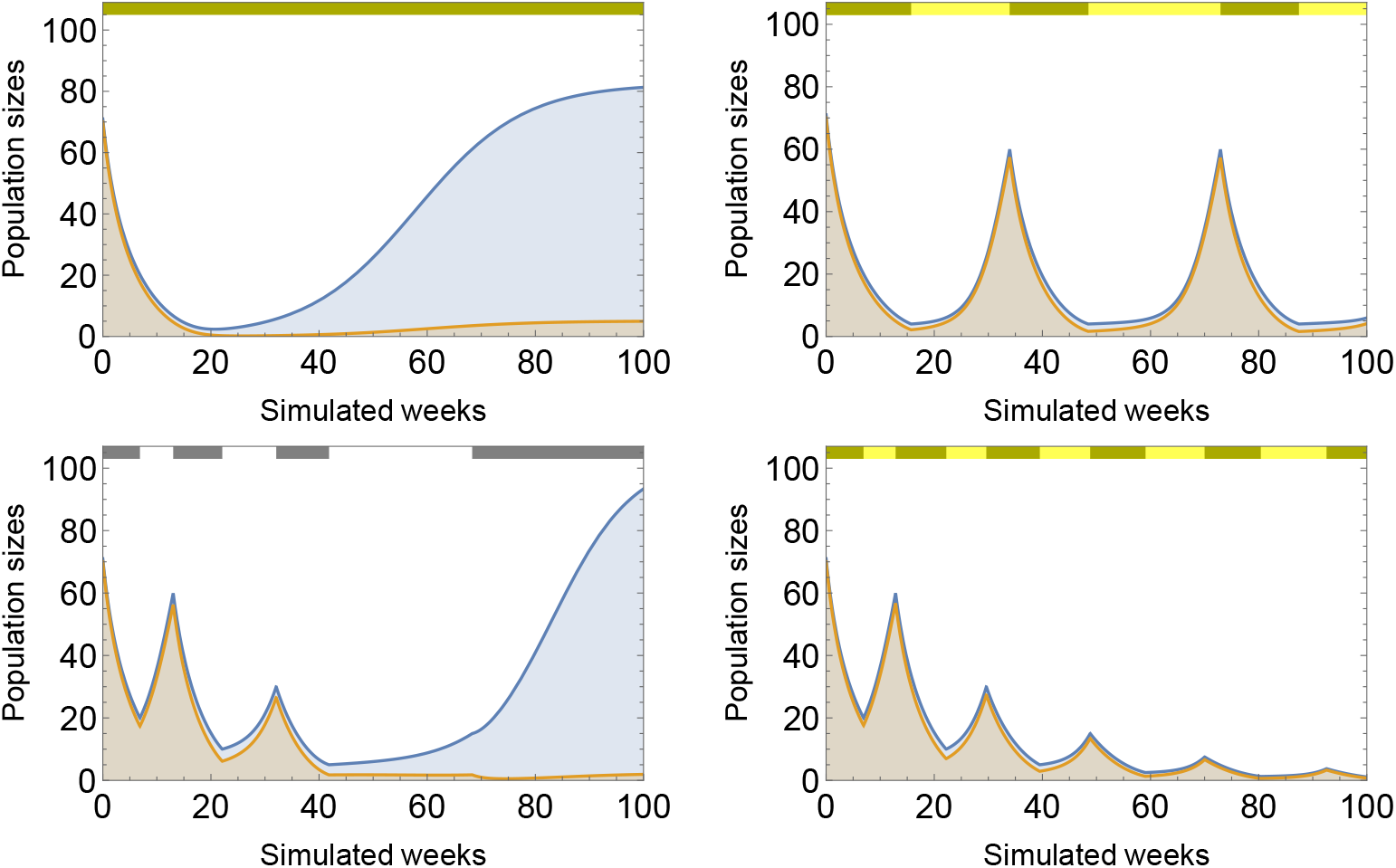
Mutation therapy when applied with cytotoxic therapy improves therapy outcomes. Continuous cytotoxic therapy combined with mutation therapy delays coordination on the resistant type (top left). Adaptive cytotoxic therapy combined with mutation therapy may reach lower composition (*g* = 32%) without losing control (top right). Therapy aimed at the extinction of the tumor fails without mutation therapy (bottom left) but succeeds with it (bottom right). Parameters: *r*_1_ = 0.4, *r*_2_ = 0.2, *d_a_* = *d_b_* = 0, *K* = 150, *c_a_* = *c_b_* = 0.05, *m_a_* = *m_b_* = 0.01, *α* = 2, *γ*_1_ = 0.4, *γ*_2_ = 0, *μ*_1_ = 0.02*, μ*_2_ = 0.

## 6 Concluding discussion

In this paper we advanced the game theory literature of cancer by proposing that coordination games, a basic class of games in non-cooperative game theory, may exist in evolutionary games of cancer. Populations playing a coordination games tend to converge on a single predominant phenotype, possibly eliminating any competing phenotypes, hence observing such an evolutionary game in its later stages obscures any alternate strategies. As a result, evolutionary coordination games may go unnoticed. We proposed a number of examples that hint at the existence of coordination games in cancer and identified the conditions under which they appear in cancer.

Coordination games in cancer may be exploited in therapy to the benefit of the patient. If the competing phenotypes react differently to therapies, the treating physician has the option to affect the balance between them. Maintaining a heterogeneous composition of two competing phenotypes within the tumor reduces its fitness. In idealized conditions this may make the difference between a successful therapy that ends with the tumor’s extinction and a failed one. In more realistic conditions adaptive therapy regimens may be designed to take advantage of coordination game by keeping the tumor’s size and composition under control, possibly indefinitely. Finally, if two competing phenotypes may be mutated into one another, new therapy regimens may be designed through which a heterogeneous composition of the tumor is maintained directly by the treating physician.

We raise four remarks, three technical and one conceptual. First, in this paper we primarily considered coordination games with two competing phenotypes. This is done primarily for expository and graphical purposes and can be generalized to include any number of phenotypes. Multi-strategy coordination games are characterized by payoff matrices where each entry of the principal diagonal is larger than any other entry in the same row or column. That is, each strategy’s fitness is maximized in its own environment, and each strategy accommodates itself more than it does any other strategy. Thus, the conditions for multi-strategy coordination games to exist become more stringent as the number of strategies increases. Each such game, however, reduced to any two of its strategies produces a two-strategy coordination game, thus identifying two-strategy coordination games is necessary for multi-strategy ones.

Secondly, our numerical examples of cancer growth and treatment all the underlying games were convex (*α* = 2). The extreme specifications are linear games (*α* = 1) in which discoordination is punished in proportion to the frequency of discoordinating players, and the extremely stringent punishments where any amount of discoordination is penalized the same way (0 payoffs if not all players agree, amounting to *α* → ∞). The former calibration would reflect the spirit of classical evolutionary games which assumes pairwise interactions between players in a well mixed population, the latter one reflects a pure coordination game played by the entirety of the population. Our calibration strikes a balance between these two extremes but our qualitative results are not sensitive to variations in *α*.

Thirdly, in our simulations we assumed a sensitive phenotype and a resistant one. This is not to suggest that phenotypes of coordination games in cancer necessarily have such a stark contrast in their response to cytotoxic therapy. Nor do we wish to make a pretense that coordination games are behind resistance mechanisms to cancer therapy. Instead, this is to showcase that, given a different response to a cytotoxic agent by two strategies of a coordination game, the physician may take advantage of the underlying game. Any difference between the response rates may be exploited to maintain a heterogeneous composition of the tumor, and our calibrations are meant to showcase the avenues of doing so. If the coordination game of cancer growth is independent of the phenotypes’ response rates to therapy, the composition may still be controlled through mutation therapy.

Finally, we raise the issue of cancer heterogeneity. It is well known that tumors display a high degree of genotypic and phenotypic heterogeneity and a rapid rate of mutation that increases heterogeneity rather than decreases it as the disease progresses. On a surface level this is in contradiction with our hypothesis that coordination games exist in cancer since they would induce a move towards homogeneity in the tumor. We argue, however, that this feature of cancer does not rule out the possibility of coordination games. The strategies of evolutionary games in cancer are generally not meant to represent a single phenotype, but usually a large number of them that coalesce into an observable trait of the tumor. We propose that coordination games take place between these traits. To continue the “rules of the road” analogy, just because the cars all drive on the right or the left side, the cars themselves may still be diverse, and even continue to branch out to display more and more heterogeneity. This is not to say that cancer’s high level of genotypic variation does not interact with the strategies of these games. In our models this interaction is captured by the rates of transmutation between the types, meaning that even in a stable, coordinated equilibrium, both types will be present.

## A Appendix: Game theory foundations

In this appendix we offer a formal discussion on the mathematical conditions of coordination games.

Evolutionary games with two strategies need to meet two sufficient and necessary conditions to produce a coordination game:

1. Each strategy proliferates at a higher rate in its own environment than in the others’ environment.
2. Neither strategy dominates the other.

Both these conditions are mild in terms of their biological implications. The former requires that the strategies be ‘selfish’, a concept that is very well documented in evolutionary biology. The latter requires that neither strategy be more fit than the other in both environments. If one strategy were to dominate the other, then the dominant strategy invariably would outcompete the dominated one.

Using the the payoff parameters *r*_1_*, r*_2_*, d*_1_*, d*_2_ as in Table 2, and assuming *r*_1_ ≥ *r*_2_ without loss of generality, (1) can be expressed as *r*_1_ ≥ *d*_1_ and *r*_2_ ≥ *d*_2_, while (2) amounts to assuming *r*_2_ ≥ *d*_1_, as *r*_1_ ≥ *d*_2_ already follows from (1).

Under these assumptions the game will have three Nash equilibria: purely type 1, purely type 2, and a mixed equilibrium where the frequency of type 1 equals *g*\*, which can be obtained by setting type 1’s payoff equal to type 2’s as follows:

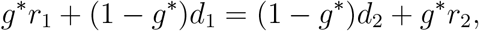

yielding 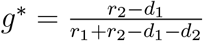, which is between 0 and 1 due to assumptions (1) and (2).

Due to (1) and (2), we have *r*_2_ ≥ *d*_1_, meaning that strategy 1 cannot invade the purely type 2 equilibrium, while the assumption *r*_1_ ≥ *r*_2_, together with (1) implies that *r*_1_ ≥ *d*_2_, hence strategy 2 also cannot invade the purely type 1 equilibrium, meaning that both pure equilibria constitute evolutionary stable strategies. On the other hand, the mixed equilibrium is unstable due to *g*\**r*_1_ + (1 − *g*\*)*d*_2_ ≤ *r*_1_ and (1 − *g*\*)*r*_2_ + *g*\**d*_1_ ≤ *r*_2_, hence both strategies are able to invade it. Under typical imitation dynamics, the mixed equilibrium separates the basins of attraction of the two pure equilibria, populations with frequencies of type 1 larger than *g*\* converge to the pure type 1 equilibrium, while those lower than *g*\* converge to the pure type 2 equilibrium.

A second avenue in which ‘coordination-like’ games may arise is through assortative matching in a game that is itself not necessarily a coordination game. In the current context, assortative matching means that a player (cancer cell) with a given strategy is more likely to interact with players of the same strategy.

For example, in a Prisoner’s Dilemma game with assortative matching, if cooperators are sufficiently more likely to interact with other cooperators, then cooperation is an evolutionary stable strategy. If, however, the assortative matching is not too strong, then desertion is also evolutionary stable.

To formalize this, take *d*_2_ > *r*_1_ > *r*_2_ > *d*_1_, which characterizes a Prisoner’s Dilemma game. In this case, neither (1) nor (2) are satisfied. Let *σ* denote the the rate of assortative matching, i.e. the probability of interaction with a player (cell) of the same type. With probability 1 − *σ*, the player interacts with an opponent chosen from a well-mixed sample of the population. Then, *σ* > (*d*_2_ − *r*_1_)/(*d*_2_ − *r*_2_) will mean that Strategy 1 (Cooperation) is evolutionarily stable as

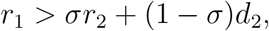

meaning that Strategy 2 (Defection) cannot invade Strategy 1.

Similarly, if *σ* < (*r*_2_ − *d*_1_)/(*r*_1_ − *d*_1_), then Type 2 (Defection) is evolutionarily stable as

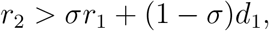

thus, Strategy 1 also cannot invade Strategy 2, producing the two pure evolutionary stable strategies of coordination games.

A *σ* that is able to satisfy both conditions exists if and only if *r*_1_ + *r*_2_ > *d*_1_ + *d*_2_. Therefore, ‘coordination-like’ games arise from the more widely studied Prisoner’s Dilemma game even if conditions (1)-(2) are not met but the sum of main-diagonal elements of the payoff matrix shown in Table 5 is larger than the sum of off-diagonal elements. The frequency of type 1 cells in the unstable mixed equilibrium that separates the basins of attraction is again denoted by *g*\*. It is calculated, as before, by setting the payoffs of the two types equal as follows:

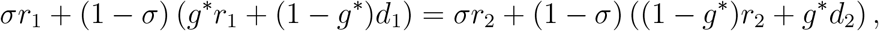

leading to

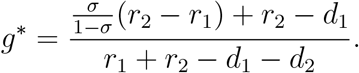

**Table 5:** An evolutionary game with two strategies is a coordination game if conditions (1) and (2) are satisfied, that is *r*_1_ ≥ *d*_1_, *r*_2_ ≥ *d*_2_, and *r*_2_ ≥ *d*_1_, with *r*_1_ ≥ *r*_2_ assumed without loss of generality. Alternatively, assortative matching can also produce ‘coordination-like’ games.

|  |  | Predominant strategy |  |
| --- | --- | --- | --- |
|  |  | Type 1 | Type 2 |
| Focal player | Type 1 | $r_1$ | $d_1$ |
| | Type 2 | $d_2$ | $r_2$ |

The main take-away from this exercise is to showcase that, although as we argue, the conditions under which coordination games can arise are mild, even when those conditions are not met, standard models of assortative matching lead to the emergence of ‘coordination-like’ games with identical transient dynamics.

